# Deep tissue infection by an invasive human fungal pathogen requires novel lipid-based suppression of the IL-17 response

**DOI:** 10.1101/2022.02.07.479457

**Authors:** P. Basso, E.V. Dang, A. Urisman, L.E. Cowen, H. D. Madhani, S. M. Noble

## Abstract

*Candida albicans* is the most common cause of human fungal infection, but the mechanisms of invasive pathogenesis remain poorly defined. Here we identify an unexpected mechanism: lipid-mediated immunosuppression. Through forward genetics, we found that *C. albicans* secretes a lipase, Lip2, that is critical for invasive disease. Murine infection with *C. albicans* strains that lack Lip2 display an exaggerated host IL-17 response that leads to fungal clearance from solid organs and host survival. IL-17 signaling is required for Lip2 action. The lipase activity of Lip2 inhibits IL-17 production indirectly through suppression of IL-23 production by tissue resident dendritic cells. We conclude that *C. albicans* suppresses antifungal IL-17 defense in solid organs by altering the tissue lipid milieu.

## Main

The yeast *Candida albicans* lives in intimate association with humans and other mammals as a beneficial component of the gut microbiota and the most common cause of fungal infectious disease^1,2^. The crucial role of the cytokine IL-17 in controlling fungal proliferation on epithelial surfaces has been exposed by studies on patients with chronic mucocutaneous candidiasis (CMC).^3^ This syndrome of persistent or recurrent *C. albicans* infections of the mouth, skin, nails, and/or vagina is linked to mutations affecting both the cytokine (IL-17A, IL-17F)^4^ and its receptor (IL-17RA, IL-17RC),^4–6^ among others. Mice bearing null alleles of the orthologous genes exhibit a similar enhanced susceptibility to mucocutaneous candidiasis^7^. On the other hand, human patients suffering from isolated CMC or treated with anti-IL-17 biologic drugs do not experience an excess risk of *C. albicans* bloodstream infections,^8^ and IL-17 is considered less important in defense against systemic fungal disease.

### *LIP2* is critical for pathogenicity during systemic infection

We identified Lip2 as a candidate fungal virulence factor in a pooled screen of ~1500 *C. albicans* GRACE (Gene Replacement And Conditional Expression) mutants^9^ in a mouse model of bloodstream infection (Basso et al., manuscript in preparation). The *lip2*^DOX-OFF^ mutant identified in the screen contains one *lip2* null allele and one *LIP2* wild-type allele that is controlled by a doxycycline (DOX)-repressible promoter (*C. albicans* is diploid). As shown in Figure 1a, *lip2*^DOX-OFF^ is strongly outcompeted by wild-type (WT) *C. albicans* in the kidneys of animals treated with doxycycline; note that kidneys are the primary target organ in the bloodstream infection model. In single strain infections, two independent *lip2^−/-^* homozygous null mutants exhibited significantly reduced lethality compared to WT or a *lip2+LIP2* strain to which a single copy of *LIP2* had been restored (Fig. 1b). These results establish that *LIP2* is required for virulence in a systemic infection model.

**Fig. 1.**
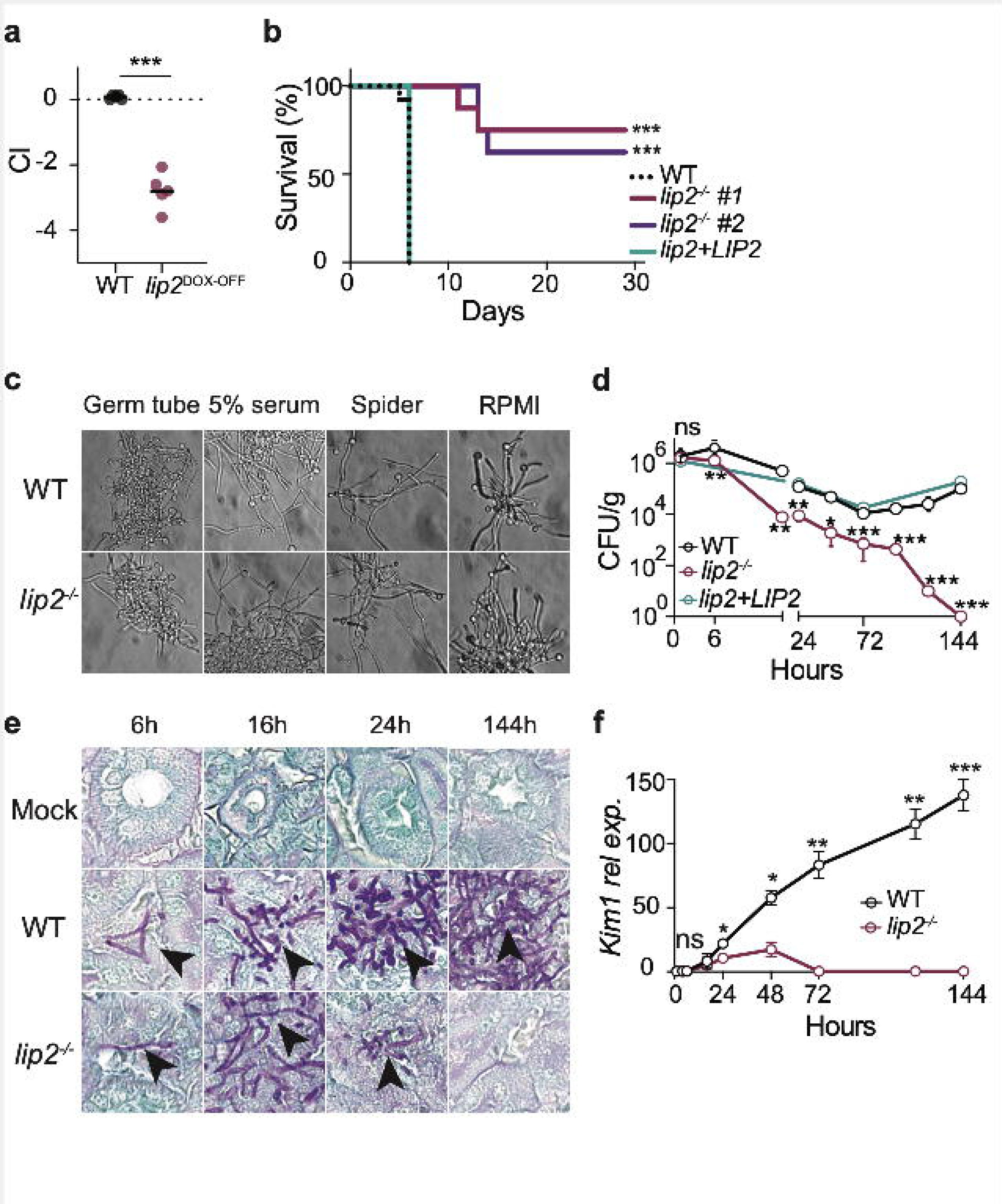
*LIP2* is required for pathogenicity. **(a)** *lip2*^DOX-OFF^ is defective for systemic virulence in the presence of doxycycline. BALB/c mice were treated with 0.25 mg/ml doxycycline via drinking water for seven days prior to retro-orbital injection with 1×10^5^ CFU of a 1:1 mixture of WT and *lip2*^DOX-OFF^; relative strain abundance in kidneys was determined using qPCR. Statistical significance was determined by a paired two-tailed t-test; ***p < 0.001. (b) *lip2^−/-^* mutants exhibit reduced lethality compared to WT or a *lip2+LIP2* gene addback strain. Groups of BALB/c mice were injected with 1×10^5^ CFU WT (n=16), *lip2*^*−*/−^ (isolate 1, n=8), *lip2*^−/-^ (isolate 2, n=8) or *lip2*^−/-^+*LIP2* (n=8). ***p < 0.001 by Mantel-Cox test. (c) *lip2*^−/-^ exhibits normal morphology under *in vitro* hypha-inducing conditions. WT and *lip2*^−/-^ were incubated under clinical “germ-tube assay” conditions (liquid YPD+5% serum), or in liquid Spider or RPMI for two hours at 37°C and 5% CO_2_. (d) *lip2*^−/-^ fails to persist in infected kidneys. Groups of BALB/c mice were infected with 1×10^5^ CFU of WT, *lip2*^−/-^, or *lip2+LIP2*, followed by euthanasia of three animals per group at the indicated time points. CFUs were determined by plating right kidney homogenates onto Sabouraud agar and counting after two days. Data represent the means and standard deviations. Statistical significance between WT and *lip2^−/-^* was determined by an unpaired two-tailed t-test. Ns nonsignificant, ** p< 0.01; ***p < 0.001. (Note that points without visible error bars displayed SEMs smaller than the circle.) (e) *lip2*^−/-^ forms normal-appearing hyphae in host kidneys. 400x image of PAS-stained left kidneys from experiment described in (D). (f) *lip2*^−/-^ causes minimal damage to kidneys. *Kim1* mRNA was measured in right kidney homogenates and normalized to *Gadph*. Statistical significance was determined by an unpaired two-tailed t-test. * p< 0.05; ** p< 0.01; ***p < 0.001; ****p< 0.0001.(Note that points without visible error bars displayed SEMs smaller than the circle.)

The best known virulence attribute of *C. albicans* is its ability to transition from budding yeast to elongated, multicellular hyphae^10^. Unlike yeast, hyphae are naturally invasive and express virulence factors such as adhesins, tissue-degrading enzymes, and the secreted toxin Candidalysin^4^. To test whether *LIP2* is required for yeast-to-hypha morphogenesis, *lip2^−/-^* and WT strains were profiled under multiple hypha-inducing conditions. Under *in vitro* conditions, *lip2^−/-^* and WT were able to filament equally well in the “germ tube” clinical diagnostic assay, as well as in YEPD+5% serum, Spider, and RPMI media maintained at 37°C (Fig.1c and Extended Data Fig. 1a). These results *s*uggest that *LIP2* is dispensable for morphogenesis. Of note, *LIP2* is itself induced under *in vitro* hypha-promoting conditions (Extended Data Fig. 1b)

To further characterize the virulence defect of the *lip2* null mutant, we performed a series of monotypic (single strain), timed infections of the bloodstream infection model with *lip2^−/-^*, WT, or saline (mock infection); a *lip2+LIP2* strain was also tested in a subset of animals. Three BALB/c mice per comparison group were euthanized after 1, 4, 6,16, 24, 48, 72, 96, 120, and 144 hours, followed by analysis of kidneys for fungal burden, pattern of fungal invasion, tissue damage, and expression of selected host transcripts. As shown in Figure 1d, *lip2^−/-^*, WT, and *lip2+LIP2* were equally represented in kidneys (~10^6^ CFU/g) after one hour of infection. Between 6 and 72 hours, the abundance of all three strains declined, but the burden of *lip2^−/-^* was reduced to a significantly greater extent. After 72 hours, the abundances of WT and *lip2+LIP2* progressively increased over the remainder of the time course, but that of *lip2^−/-^* continued to decline until it became undetectable at 144 hours. Fungal titers were also determined in the liver and spleen at selected time points (Extended Data Fig. 1c). All three fungal strains were similarly abundant (~10^3^ CFU/g) in both organs after 1 hour. In the liver, WT and *lip2+LIP2* remained relatively stable, but *lip2^−/-^* became undetectable by 144 hours. All three strains were rapidly cleared from the spleen. This analysis indicates that *LIP2* is not necessary for the initial entry of *C. albicans* into solid organs, but it is required to maintain a stable infection in the kidney and liver.

Patterns of fungal invasion were assessed in sectioned left kidneys after staining with Periodic Acid Schiff (PAS). As shown in Figures 1e and Extended Data Fig. 1d, *lip2^−/-^* formed normal-appearing hyphae in kidneys, suggesting that *lip2^−/-^* is competent for morphogenesis in the host as well as under *in vitro* conditions (Fig. 1c). However, stark differences were apparent in the extent of renal invasion by *lip2^−/-^* vs. WT. By 16 hours, clusters of WT hyphae could be visualized throughout the renal cortex (Extended Data Fig. 1e), a highly perfused region that receives ~25% of cardiac output^11,12^. Over the remaining time course, the areas occupied by WT progressively enlarged and coalesced until, by 144 hours, they spanned the renal cortex, corticomedullary junction, and portions of the medulla. By contrast, *lip2^−/-^* formed much smaller collections at 16 hours (fig. S1e), and these diminished in size over subsequent time points, eventually becoming virtually undetectable at 144 hours (Extended Data Figs. 1 d & e).

Kidney damage was evaluated histologically on hematoxylin and eosin (H&E)-stained left kidney sections. Starting at 16 hours, WT-infected kidneys exhibited areas of tissue necrosis with microabscess formation at the sites of hyphal infiltration (Extended Data Fig. 1f). These areas of injury expanded and progressed over the subsequent time course, resulting in interstitial edema, widespread acute tubular injury and parenchymal necrosis, and frank hemorrhage. By contrast, *lip2^−/-^-*infected organs remained free of significant injury over the time course, despite the presence of localized microabscesses that peaked at 24 hours and subsequently resolved (Extended Data Fig. 1f). We next used RT-qPCR to evaluate right kidney homogenates for *Kim1* (Kidney Injury Molecule-1) mRNA, a highly sensitive marker of acute renal injury^13^. As shown in Figure 1f, small increases of *Kim1* were apparent after 24 and 48 hours of infection with *lip2^−/-^*, followed by a return to baseline (Fig. 1f). By comparison, WT-infected kidneys exhibited significantly larger elevations of *Kim1* at 24 and 48 hours, with progressive increases continuing over the remainder of the experiment (Fig. 1f). These results indicate that, consistent with its reduced titers after ~24 hours, *lip2^−/-^* causes only minor, transient injury to the host.

### IL-17 triggers an antifungal response during invasive candidiasis

We considered two potential explanations for the persistence defect of *lip2^−/-^*: The encoded lipase may be required for normal proliferation in tissues, or it may defend against clearance by the host. To gauge the host immune response to *lip2^−/-^* versus WT strains in infected kidneys, RT-qPCR was used to quantify RNA expression for six proinflammatory cytokines in kidney homogenates. As shown in Figure 2a, expression of *Il1β, Il6, Il10*, and *Tnf* was significantly higher in kidneys infected with WT compared to *lip2^−/-^*, whereas *Ifng* was minimally induced by either strain. *Il17a* was the only cytokine to be expressed more strongly in *lip2-*infected organs, as early as 6 hours after infection. The latter result was confirmed in additional animals, where infection with the *lip2^−/-^* mutant was found to be associated with enhanced expression of *Il17a* mRNA and, following a delay, IL-17A protein (Fig. 2b).

**Fig. 2.**
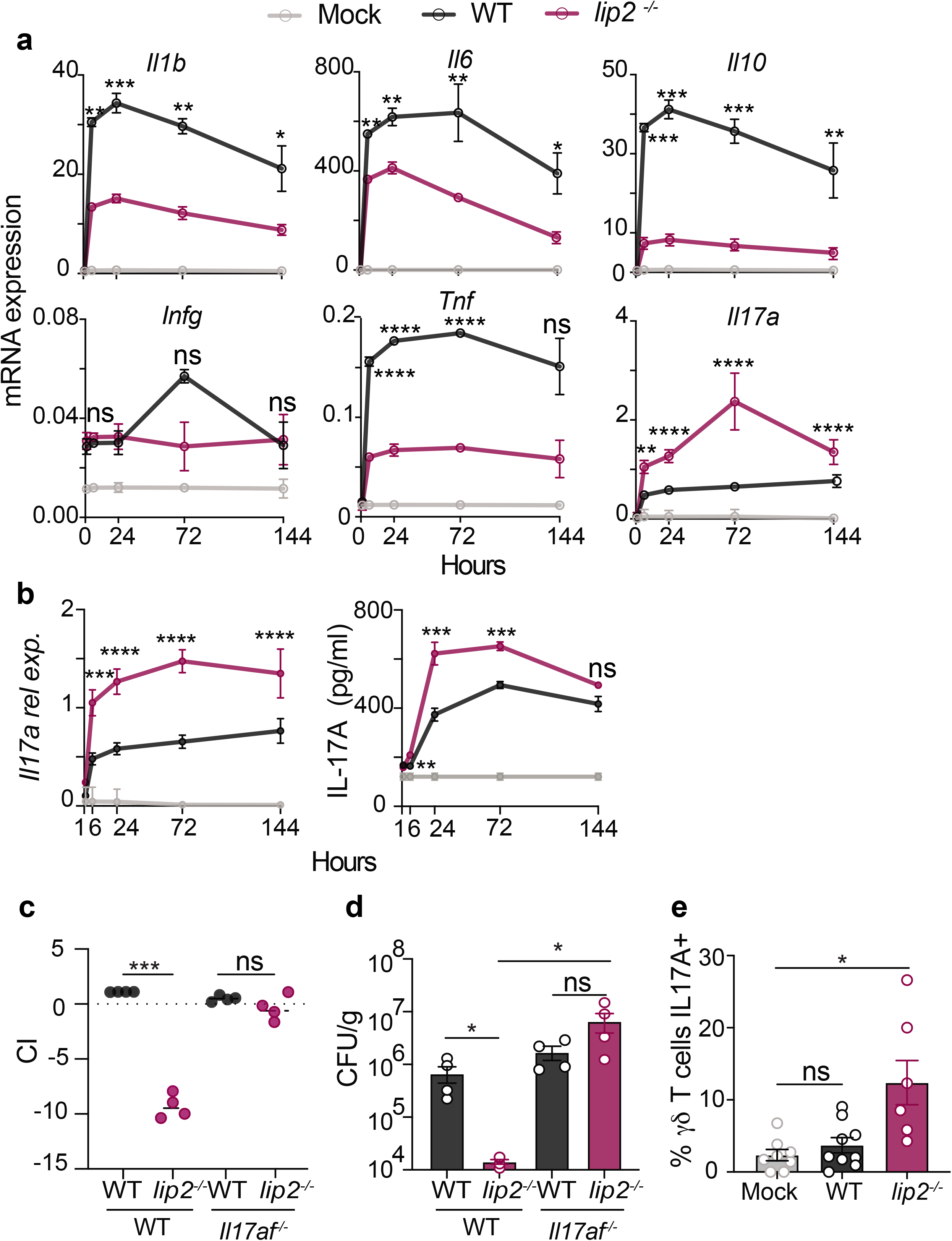
IL-17 mediates an antifungal response during systemic infection. **(a)** *lip2* provokes an exaggerated *Il17a* response in infected kidneys. Groups of BALB/c mice were infected with 1×10^5^ CFU of WT or *lip2*^−/-^ followed by euthanasia of three animals per group at the indicated time points. *Il1b*, *Il6*, *Il10*, *Ifng, Tnf*, *Il17a* mRNA expression was assessed by RT-qPCR in kidney homogenates at the indicated time points (expression relative to *Gadph*). Statistical significance between WT and *lip2*^−/-^ was determined by an unpaired two-tailed t-test. * p< 0.05; ** p< 0.01; ***p < 0.001; ****p< 0.0001. (Note that points without visible error bars displayed SEMs smaller than the circle.) (b) *lip2^−/-^* induces a strong renal IL-17A response during systemic infection. Groups of BALB/c mice were infected with 1×10^5^ CFU WT or *lip2*^*−*/−^ followed by euthanasia of three animals per group at the indicated time points. *Il17a* mRNA was measured by RT-qPCR in right kidney homogenates and normalized to *Gadph* (left panel). IL-17A protein production was evaluated by ELISA in the left kidney homogenates (right panel). Statistical significance between WT and *lip2*^−/-^ was determined by an unpaired two-tailed t-test. ns: p=0.1797; ** p< 0.01; ***p < 0.001; ****p< 0.0001. (c) The *lip2*^−/-^ defect of fitness is suppressed in *Il17af^−/-^* mice. Groups of four WT or *Il17af^−/-^* C57BL/6J mice were infected with 1×10^5^ CFU of a 1:1 mixture of WT and *lip2*^−/-^; Mice were euthanized after seven days and relative strain abundance in kidneys was determined using qPCR. Statistical significance was determined by a paired two-tailed t-test; ns: p= 0.0685; ***p < 0.001. (d) *lip2^−/-^* clearance from kidneys requires IL-17. Groups of four WT or *Il17af^−/-^* C57BL/6J mice were infected with with 5×10^5^ CFU of WT or *lip2*^−/-^ strains. Animals were euthanized after four days and the fungal burden in kidneys was determined by plating right kidney homogenates onto Sabouraud agar. Statistical significance was determined by one-way ANOVA (Turkey’s multiple comparisons test); ns: p= 0.0988; *p < 0.05 (e) Renal **γ**δ T cells produce IL-17A upon *lip2*^−/-^ stimulation. Groups of C57BL/6J mice were infected with with 1×10^6^ CFU of WT or *lip2*^−/-^. Animals were euthanized after six hours of infection. Flow cytometry analysis was performed on total renal cells after four hours of stimulation with PMA, ionomycin and Golgi STOP (Note: see Extended Data Fig. 2 for FACS gating). Statistical significance was determined by one-way ANOVA (Turkey’s multiple comparisons test); ns: p= 0.8339; **p < 0.01.

We reasoned that, if IL-17 plays a role in eliminating *lip2^−/-^* from infected kidneys, then the virulence defect of the mutant would be suppressed in animals that do not produce IL-17. Consistent with this prediction, *lip2^−/-^* exhibits normal competitive fitness against WT in *Il17af^−/-^* mice, but not in *Il17af^+/+^* control animals (Fig. 2c). Of note, infection of C57B/6J mice (the genetic background of *Il17af^−/-^* animals) with the *lip2^−/-^* mutant produced similar changes in renal *Il17a* and IL-17A to those observed in BALB/c mice (Fig. 2b and Extended Data Fig. 2a). Fungal persistence was also compared in *Il17af^−/-^* and *Il17af^+/+^* animals. After 4 days of systemic infection, there was no significant difference between the abundances of *lip2^−/-^* and WT strains in kidneys of *Il17af^−/-^* animals, whereas *lip2^−/-^* was significantly depleted in kidneys of *Il17af*^+/+^ animals (Fig. 2d). These results suggest that the virulence defect of *lip2^−/-^* likely results from an enhanced IL-17 response. Moreover, the finding that IL-17 is required to eliminate *lip2^−/-^* from kidneys implies that IL-17 directs potent antifungal activity during systemic candidiasis.

To probe the mechanism by which Lip2 might modulate host immunity, we began by asking which cells produce IL-17 in infected kidneys (Fig. 1f). Th17 cells are generally considered to be a major source of IL-17 in mammals^3,8^, and this cell type has previously been shown to protect against systemic candidiasis in mice that are also intestinally colonized with *C. albicans*^14^. However, CD4^+^ T cells are poorly represented in mouse kidneys over the first 24 hours of *C. albicans* bloodstream infection^12^, whereas renal *Il17a* mRNA increases within 6 hours (Fig. 2a, 2b). We therefore focused on tissue-resident immune cells that are capable of producing IL-17, such as natural killer T cells (NKT), innate lymphoid cells (ILC), and **γ**δ T cells; note that **γ**δ T cells have previously been reported to produce IL-17 in kidneys during systemic candidiasis^15^. Kidneys were recovered from C57B/6J mice 6 or 24 hours after inoculation with the *lip2*^−/-^ mutant, WT, or normal saline (“mock” infection), followed by cell dissociation and staining for intracellular IL-17A and leukocyte surface markers. As shown in Figure 2e and Extended Data Fig. 2b, **γ**δ T cells (but not ILC3s or CD4+ T cells) were identified as a source of IL-17 in *lip2*^−/-^*-*infected kidneys. Similar results were obtained using SMART17A mice (in the C57B/6J background) that display a human low-affinity nerve growth factor receptor marker on the surface of cells that express mouse *Il17a* (Extended Data Fig. 2c)^16^.

### Lip2 suppresses the activation of renal dendritic cells

Because **γ**δ T cells typically upregulate IL-17 in response to an upstream cytokine, such as IL-23^8^, we investigated *Il23* expression in infected kidneys. As shown in Figure 3a, renal *Il23* mRNA was significantly elevated within 6 hours of infection with WT *C. albicans*, and the effect was even stronger in *lip2*^−/-^-infected organs. A similar pattern is observed with IL-23 protein, following a delay (Fig. 3a). Macrophages and dendritic cells (DC) are common sources of IL-23 in solid organs. Using flow cytometry, DCs and macrophages were quantified in kidneys recovered after 6 hours of infection with WT or *lip2*^−/-^, or in mock-infected mice. As shown in Extended Data Fig. 3a, 3b and 3c, DCs composed ~15% of total renal CD45^+^ cells, regardless of infection with *C. albicans*, whereas the macrophage population was too small to measure with accuracy. Using RT-qPCR to quantify *Il23* expression in isolated renal DCs, we observed significant upregulation after 6 hours of infection with WT *C. albicans* and an even stronger response to the *lip2^−/-^* strain (Fig. 3b). In support of DC activation by *C. albicans*, flow cytometry revealed stepwise increases in surface expression of MCHII in DCs from in organs infected with WT or *lip2^−/-^* (Fig. 3c). Together, these results suggest that tissue-resident DCs respond to fungal invasion of the kidney by producing IL-23, a known activator of IL-17 expression. For reasons that remain unclear, this response is exaggerated in the presence of *lip2^−/-^*.

**Fig. 3.**
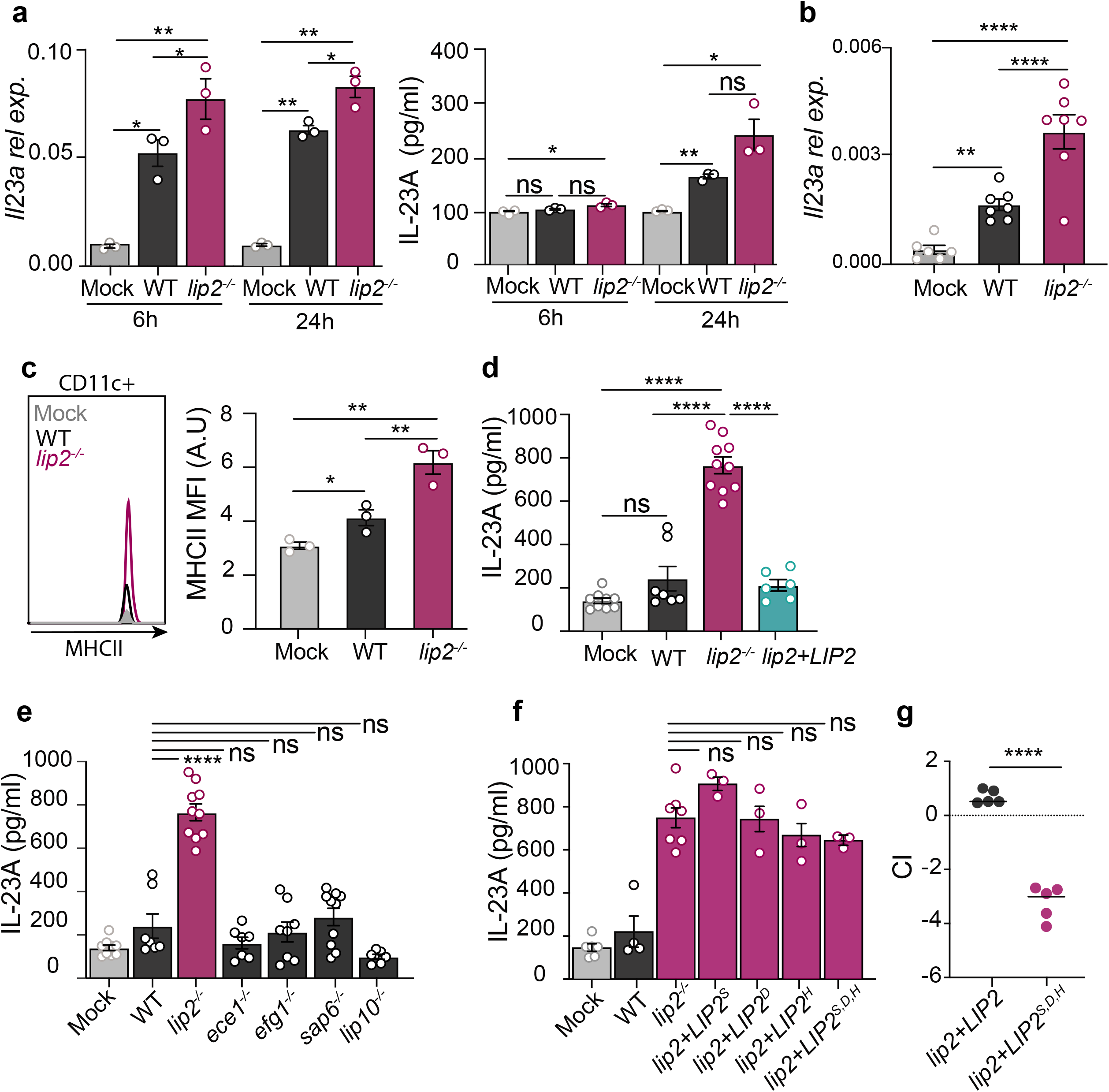
Lip2 suppresses the activation of renal dendritic cells. **(a)** *lip2*^−/-^ induces IL-23A production in kidneys. Groups of C57BL/6J mice were infected with 1×10^6^ CFU of WT or *lip2*^−/-^ followed by euthanasia of three animals per group at the indicated time points. *Il23a* mRNA was measured by RT-qPCR in right kidney homogenates and normalized to *Gadph* (left panel). IL-23A protein production was evaluated by ELISA in the left kidney homogenates (right panel). Statistical significance was determined by one-way ANOVA (Tukey’s multiple comparisons test); * p< 0.05; ** p< 0.01. (b) Kidney resident DCs respond to *C. albicans* invasion. Groups C57BL/6J mice were infected 1×10^6^ CFU of WT or *lip2*^−/-^ strains for six hours. Renal DCs were isolated and *Il23a* mRNA levels were measured by RT-qPCR and normalized to *Gadph*. Statistical significance was determined by one-way ANOVA (Turkey’s multiple comparisons test); ** p< 0.01; *** p<0.001; **** p<0.0001. (c) *LIP2* is required to prevent cell surface expression of MCHII by renal DCs in infected kidneys. Groups C57BL/6J mice were infected 1×10^6^ CFU of WT or *lip2*^−/-^ for six hours. DCs were analyzed by flow cytometry for MCHII. Statistical significance was determined by one-way ANOVA (Tukey’s multiple comparisons test); * p<0.05; ** p< 0.01. (d) Lip2 suppresses IL-23 production by BMDCs. BMDCs from C57BL/6J mice were cocultured with WT, *lip2*^−/-^, or *lip2+LIP2* at an MOI of 1 for two hours followed by measurement of IL-23A in cell supernatants by ELISA. Statistical significance was determined by one-way ANOVA (Tukey’s multiple comparisons test); ns: p= 0.6247; **** p<0.0001. (e) BMDC activation is specific to *lip2*. Coculture experiment with BMDC and *C. albicans* virulence-defective mutants (*efg1^−/-^, ece1^−/-^, sap6^−/-^*) and paralog of *lip2^−/-^* (*lip10^−/-^*) was performed as described in (D). (f) Lip2 lipase activity is required to prevent the activation of BMDCs. Coculture experiment with BMDC and *C. albicans* strains expressing Lip2 point mutants affecting conserved residues at the catalytic site (*lip2*+*LIP2^S196A^, lip2*+*LIP2^D240A^, lip2*+*LIP2^H344A^, lip2*+*LIP2^S196A,D240A,H344A^)* was performed as described in (D). (g) Lip2 lipase activity is required for virulence. BALB/c mice were infected with 1×10^5^ CFU of a 1:1 mixture of *lip2*+*LIP2* and *lip2*+*LIP2^S196A,D240A,H344A^*. The relative strain abundance in kidneys was determined by qPCR. Statistical significance was determined by a paired two-tailed t-test; ****p < 0.0001.

To test whether *C. albicans* can directly activate IL-23 production by DCs, coculture experiments were performed with bone marrow-derived dendritic cells (BMDC) prepared from C57B/6J mice. BMDCs were incubated at a MOI of 1 with WT, *lip2^−/-^*, three additional virulence-defective mutants, a *lip10^−/-^* mutant, or cell medium alone for 2 hours, followed by quantitation of IL-23 in culture supernatants. As shown in Figure 3d, exposure to the *lip2^−/-^* strain but not WT induced a strong IL-23 response from cocultured BMDCs. The inducing activity was specific to *lip2^−/-^*, as no induction was observed by exposure to mutants defective in Candidalysin production (*ece1*)^17^, yeast-to-hypha morphogenesis (*efg1^−/-^*)^18^, a secreted aspartyl protease (*sap6^−/-^*)^19^, or the closest paralog of Lip2 (*lip10^−/-^*) (Fig. 3e). These results suggest that Lip2 may function to suppress IL-23 production by DCs.

### The lipase activity is crucial for the immunomodulatory action

*LIP2* encodes a secreted lipase with nine paralogs in *C. albicans*^20^. To determine whether lipase activity is required for the immune suppressive effect of Lip2, we engineered four *LIP2* point mutants (*lip2*^S196A^, *lip2*^D240A^, *lip2*^H344A^, and *lip2*^S196A,D240A,H344A^), targeting conserved residues in the predicted lipase domain (Extended Data Fig. 4a). FLAG-tagged versions of each mutant allele and wild-type *LIP2* were individually introduced into a *lip2^−/-^* null mutant, and immunoblotting of cell supernatants was performed to verify that each of the proteins was expressed (Extended Data Fig. 4a). Culture supernatants were then assessed for *in vitro* lipase activity. As shown in figure S4B, each point-mutant strain exhibited significantly less lipase activity than *lip2+LIP2* or WT *C. albicans*, with similar activity to the *lip2^−/-^* null mutant, suggesting that the encoded proteins are catalytically inactive. Moreover, all four mutants induced high levels of IL-23 expression following 2 hours of coculture with BMDCs (Fig. 3f), similar to the phenotype of the *lip2^−/-^* null mutant. Finally, we tested the fitness of the *lip2+LIP2*^S196A^, *lip2+LIP2*^D240A^, and lip2+*LIP2*^S196A,D240A,H344A^ strains in 1:1 competition with WT in the mouse systemic infection model. All three tested mutants exhibited reduced virulence (Fig. 3g and Extended Data Fig. 4c), similar to the phenotype of *lip2^−/-^*. Together, these experiments suggest that Lip2 lipase activity is required for suppression of the BMDC response to *C. albicans* and for virulence in the host.

### Palmitic acid suppresses the activation of the dendritic cells

Lipases catalyze the hydrolysis of triglycerides to fatty acids and glycerol. We hypothesized that, within the host, Lip2 may modulate DC reactivity by increasing the local concentration of one or more fatty acids. To test this hypothesis, we assayed the ability of several common fatty acids to suppress the induced IL-23 response of BMDCs to *lip2^−/-^*. As shown in Figure 4a and Extended Data Fig. 4d, 4a, 0.1 μM palmitic acid but not stearic acid or linoleic acid suppressed IL-23 production by BMDCs exposed to *lip2^−/-^* to the level observed in unexposed BMDCs (or WT-exposed BMDCs). Likewise, 0.1 μM palmitic acid suppressed the response of BMDCs to strains expressing catalytically inactive alleles of *LIP2* (Extended Data Fig. 4f). We compared the ability of WT, *lip2^−/-^*, *lip2+LIP2*, and strains expressing *LIP2* point mutants to hydrolyze a pNp-palmitate substrate. As shown in Figure 4b, the WT and *lip2+LIP2* strains exhibited significantly greater activity than the null mutant or point mutant strains. Thus, Lip2 is capable of liberating palmitate, a fatty acid that suppresses IL-23 production by BMDCs during coculture with *C. albicans*.

**Fig. 4.**
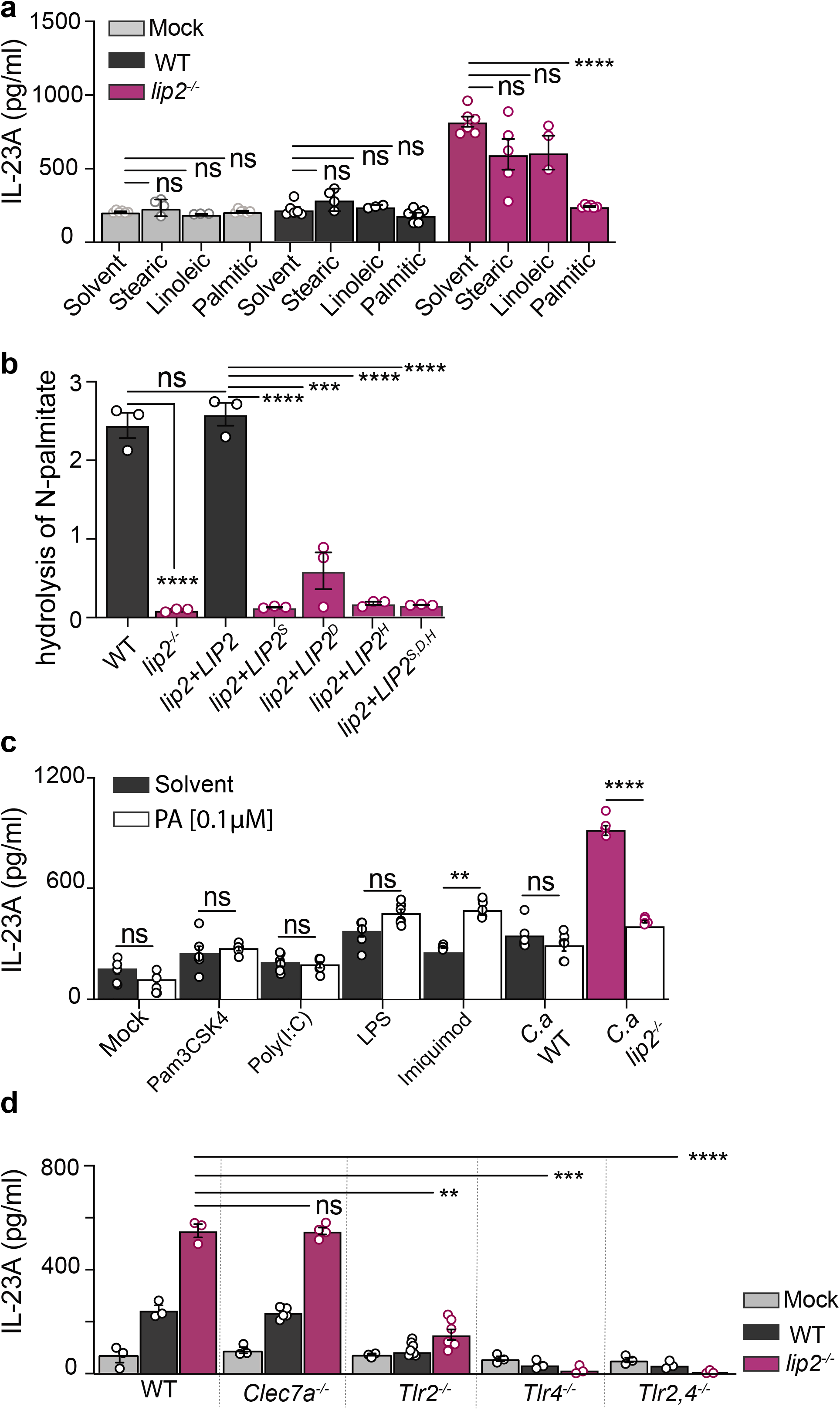
Predicted catalytic residues of Lip2 is required for its immunomodulatory role. **(a)** Palmitic acid suppresses the activation of BMDCs by *lip2^−/-^*. BMDCs from C57BL/6J mice were incubated with 0.1 μM of fatty acids (palmitic acid, stearic acid or linoleic acid) solubilized in chloroform (chloroform (solvent) was used as control) and co-cultured with WT, *lip2*^−/-^ strain at a MOI of 1 for two hours followed by measurement of IL-23A in cell supernatants by ELISA. Statistical significance was determined by one-way ANOVA (Turkey’s multiple comparisons test); ns: p> 0.05; **** p<0.0001. (b) Lip2 releases palmitic acid. pNp-palmitate hydrolysis was measured after an incubation of pNp-palmitate with the WT, *lip2^−/-^*, *lip2+LIP2* and lip2 catalytic mutants supernatant from liquid culture at OD_600_=1. Statistical significance was determined by one-way ANOVA (Turkey’s multiple comparisons test); ns: p> 0.05; *** p<0.001; **** p<0.0001. (c) The dampening effect of palmitic acid is specific to the *lip2^−/-^* stimulus. BMDCs from C57BL/ 6J mice were incubated with 0.1 μM of chloroform (solid bars) or with 0.1 μM of palmitic acid solubilized in chloroform (clear bars). BMDCs were stimulated for two hours with TLR ligands (Pam3CSK4: TLR1/2; Poly(I:C): TLR3; LPS: TLR4; imiquimod: TLR7/8) or cocultured with WT, *lip2*^−/-^ strain at a MOI of 1 followed by measurement of IL-23A in cell supernatants by ELISA. Statistical significance was determined by an unpaired two-tailed t-test; ns: p> 0.05; ** p<0.001; **** p<0.0001. (d) TLR2 and TLR4 are required for the BMDCs activation. BMDCs from C57BL/6J mice were incubated with 0.1 μM of fatty acids (palmitic acid, stearic acid or linoleic acid) solubilized in chloroform (chloroform (solvent) was used as control) and co-cultured with WT, *lip2*^−/-^ strain at an MOI of 1 for two hours followed by measurement of IL-23A in cell supernatants by ELISA. Statistical significance was determined by one-way ANOVA (Turkey’s multiple comparisons test); ns: p> 0.05; **** p<0.0001.

The immune dampening effect of palmitic acid on DC activation by *C. albicans* was somewhat surprising in light of published reports on the proinflammatory activity of palmitic acid on macrophages and DCs, particularly in the context of chronic diseases such as obesity and type 2 diabetes mellitus^21^. Especially relevant is a study showing that palmitic acid strongly enhances the response of BMDCs to a panel of TLR agonists^22^. The authors of this study found that, following a 24-hour incubation, 500 **μ**M palmitic acid reduces the levels of IL-23 mRNA and protein produced by BMDCs exposed to Pam3CSK4 (TLR1/TLR2 agonist), poly(I:C) (TLR3 homodimer agonist), LPS (TLR4 homodimer agonist), or imiquimod (TLR7/TLR8 agonist). To clarify whether palmitic acid exerts distinct anti-inflammatory versus pro-inflammatory effects on BMDCs exposed to different stimuli, we measured IL-23 expression upon exposure to *lip2^−/-^* under our assay conditions (2 hours, 0.1 μM palmitic acid) and the previously reported conditions (24 hours, 500 μM palmitic acid), using Pam3CSK4, poly(I:C), LPS, and imiquimod as controls. As shown in Figure 4c, 0.1 μM palmitic acid significantly enhanced the IL-23 response to imiquimod but not to the other TLR ligands under our assay conditions, but suppressed the response to *lip2^−/-^*. Under the previously reported conditions, 500 μM palmitic acid significantly enhanced the IL-23 response to LPS and imiquimod but suppressed the response to *lip2^−/-^* (Extended Data Fig. 4g). These results suggest that the immune modulatory activity of palmitic acid on BMDCs varies with the identity of the stimulus (LPS and imiquimod vs. the *lip2^−/-^* strain).

TLR2, TLR4, and Dectin-1 are cell-surface associated pattern recognition receptors that have previously been implicated in immune recognition of *C. albicans*^23^. TLR2 and TLR4 have been suggested to recognize O-linked mannose residues while Dectin-1 recognizes β-1,3-linked glycan residues of the fungal cell wall. To determine whether any of these receptors plays a nonredundant role in the IL-23 response to *lip2^−/-^*, we performed coculture assays with *C. albicans* and BMDCs prepared from *clec7a^−/-^* (Dectin-1 locus), *tlr2^−/-^, tlr4^−/-^*, or *tlr2,tlr4^−/-^* mice. As shown in Figure 4d, *Clec7a* is dispensable for BMDC activation by *lip2^−/-^*, but the IL-23 response is significantly diminished in cells lacking *Tlr2* or *Tlr4* and virtually eliminated in cells lacking both *Tlr2* and *Tlr4*.

Our results support a model in which Lip2 promotes fungal virulence by suppressing the host IL-17 response. We propose that, in organs such as the kidney, Lip2 increases the local concentration of immune modulatory fatty acids such as palmitic acid, thereby blunting TLR2- and TLR4-dependent activation of tissue resident DCs. The immune dampening effect may be mediated directly on DCs or it may occur indirectly, for example because of decreased expression of TLR2/TLR4 ligands on the *C. albicans* cell surface. In contrast, during infections with *C. albicans* mutants that fail to express *LIP2* or that express a catalytically inactive version of the enzyme, DCs are activated to release IL-23, leading to IL-17 expression by tissue resident **γ**δ T cells. The lipid-mediated suppression of IL-17 that we observe in solid organs is apparently not active in skin^24^. This difference may be related to the observation that, in skin, *C. albicans* is sensed by nociceptive sensory fibers, which leads to DC activation and IL-17 production by dermal **γ**δ T cells^24^. This neuro-immune pathway may bypass Lip2-mediated DC suppression that we observe in solid organs.

## Acknowledgments

We are grateful to Prof. Gregory Barton (UC Berkeley) for the gift of *tlr2^−/-^, tlr4^−/-^* and *tlr2,4^−/-^* femurs and to Prof. Richard Locksley (UCSF) for the gift of SMART17 mice. We thank Allison Cohen for technical advice on the preparation of BMDCs. Finally, we are endebted to Prof. Anita Sil (UCSF), Prof. Sarah L. Gaffen (U. Pittsburg), and Prof. Partha S. Biswas (U. Pittsburg) for illuminating discussions and advice. We thank Merck and Genome Canada for making the original *C. albicans* GRACE mutant collections available.

## Author contributions

Conceptualization: PB, SMN

Methodology: PB, EVD, SMN

Investigation: PB, EVD

Visualization: PB, AU, SMN

Funding acquisition: HDM, LEC, SMN

Project administration: SMN

Supervision: HDM, LEC, SMN

Writing – original draft: PB, SNM

Writing – review & editing: PB, SNM, EVD, HDM

## Data and materials availability

“All data are available in the main text or the Extended Data materials.”

---

**Extended Data Fig. 1.**
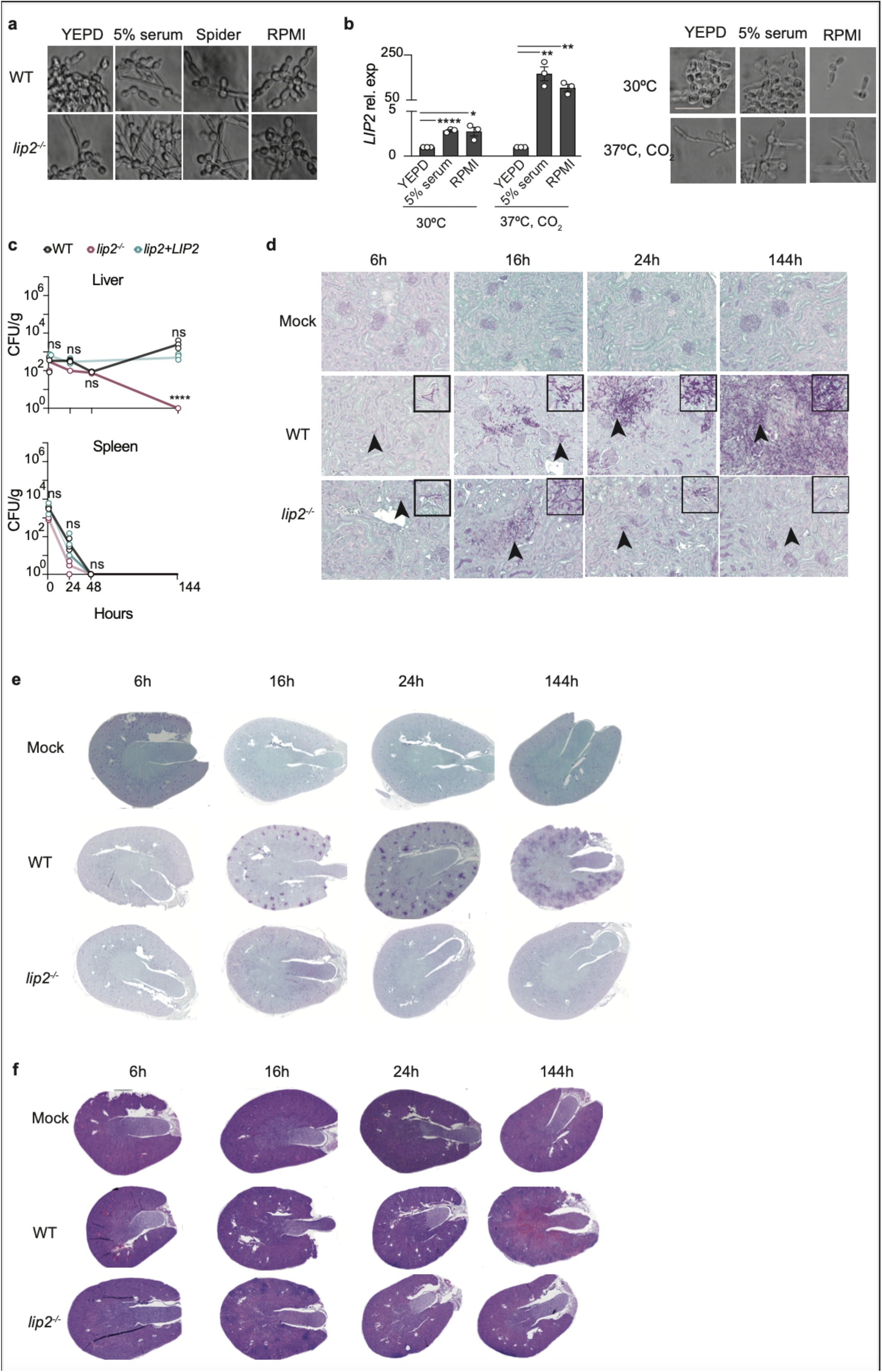
**(a)** *lip2^−/-^* exhibits normal morphology under in vitro liquid conditions. WT and *lip2^−/-^* strains were incubated in YPD, YPD+5% serum, Spider and RPMI for two hours at 30C. (b) *LIP2* is expressed under in vitro hypha-promoting conditions. *LIP2* mRNA was measured in cultures grown to OD_600_=1 in the indicated media. mRNA expression was normalized to *ACT1*. Pictures of the morphology of WT and *lip2^−/-^* strain in indicated conditions were taken at OD_600_=1. Statistical significance was determined by an unpaired two-tailed t-test; * p<0.05; ** p<0.001; **** p<0.0001 (c) Lip2 is required to persist in liver. Groups of BALB/c mice were infected with 1×10^5^ CFU of WT and *lip2^−/-^* and *lip2+LIP2* followed by euthanasia of three animals per groups at the indicated time points (note that the CFU from liver and spleen were from the same animals as the CFU from kidneys presented in Fig.1d). The numbers of recovered CFU from organs are indicated in the plots for individual animal. Statistical significance determined by an unpaired two-tailed t-test. ns: p> 0.05. (d) *lip2^−/-^* forms small micro abscesses before its clearance. 400X images of PAS-stained left kidneys from experiment described in Fig.d and presented in Fig.1e. (e) Lip2 is required to invade the renal medulla. 40X images of PAS-stained of the left kidney from experiment described in Fig.1d. (f) Kidney damages are visualized in WT-infected conditions. 40X images of H&E-stained of the left kidney from experiment described in Fig.1D

**Extended Data Fig. 2.**
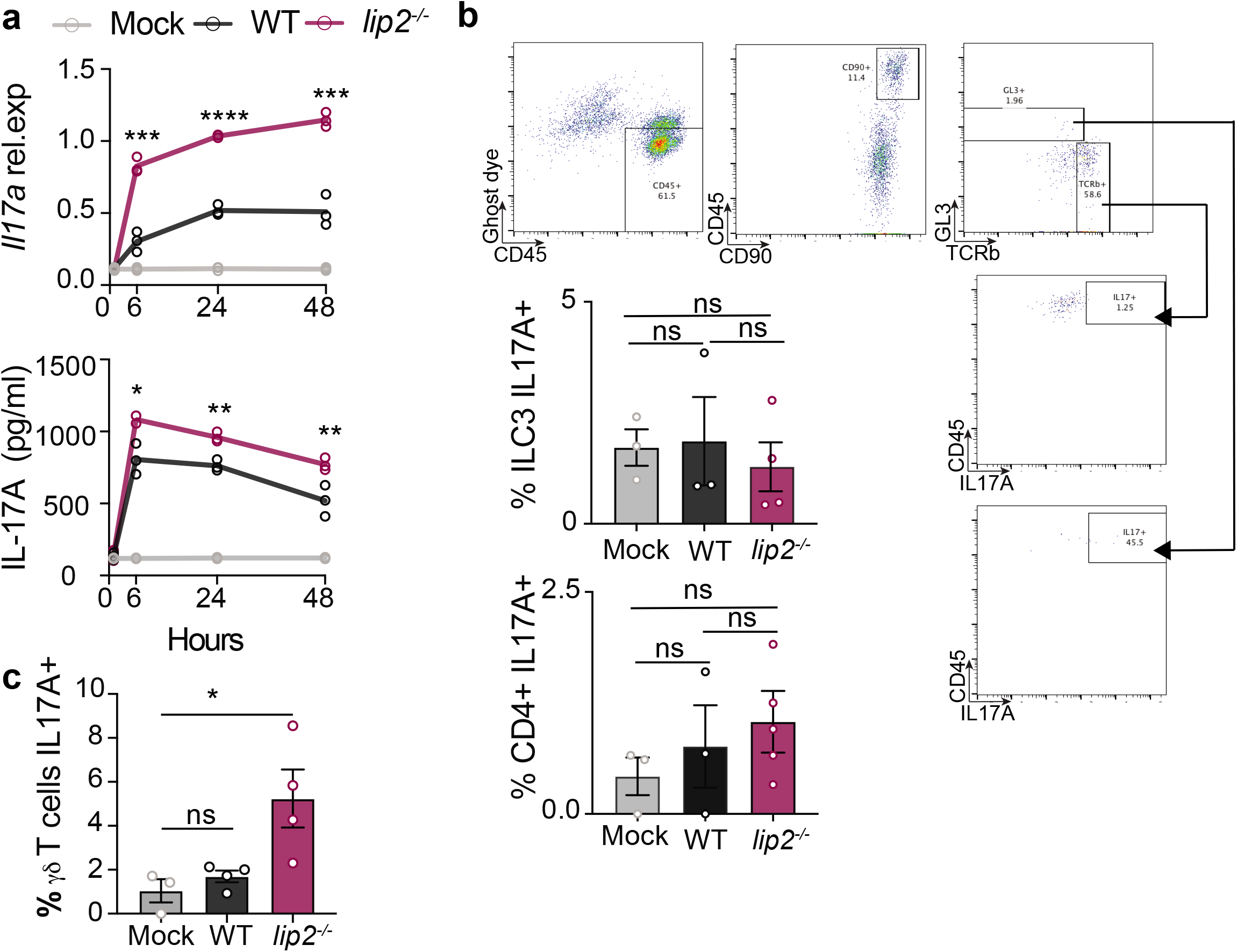
**(a)** *lip2^−/-^* induces a strong renal IL-17A response during systemic infection. Groups of C57BL/6J mice were infected with 1×10^6^ CFU WT or *lip2*^*−*/−^ followed by euthanasia of three animals per group at the indicated time points. *Il17a* mRNA was measured by RT-qPCR in right kidney homogenates and normalized to *Gadph* (upper panel). IL-17A protein production was evaluated by ELISA in the left kidney homogenates (lower panel). Statistical significance between WT and *lip2*^−/-^ was determined by an unpaired two-tailed t-test. ** p< 0.01; ***p < 0.001; ****p< 0.0001. (b) Renal **γ**δ T cells produce IL-17A upon *lip2*^−/-^ stimulation. Groups of C57BL/6J mice were infected with with 1×10^6^ CFU of WT or *lip2*^−/-^ strains. Animals were euthanized after six hours of infection. Flow cytometry analysis was performed on total renal cells after four hours of stimulation with PMA, ionomycin and Golgi STOP (Note: see Fig. 2E for quantification). Statistical significance was determined by one-way ANOVA (Turkey’s multiple comparisons test); ns: p>0.05. (c) Renal **γ**δ T cells produce IL-17A upon *lip2*^−/-^ stimulation. Groups of C57BL/6J SMART17 mice were infected with with 1×10^6^ CFU of WT or *lip2*^−/-^ strains. Animals were euthanized after six hours of infection. Flow cytometry analysis was performed on total renal cells. Statistical significance was determined by one-way ANOVA (Turkey’s multiple comparisons test); ns: p>0.05; * p<0.05.

**Extended Data Fig. 3.**
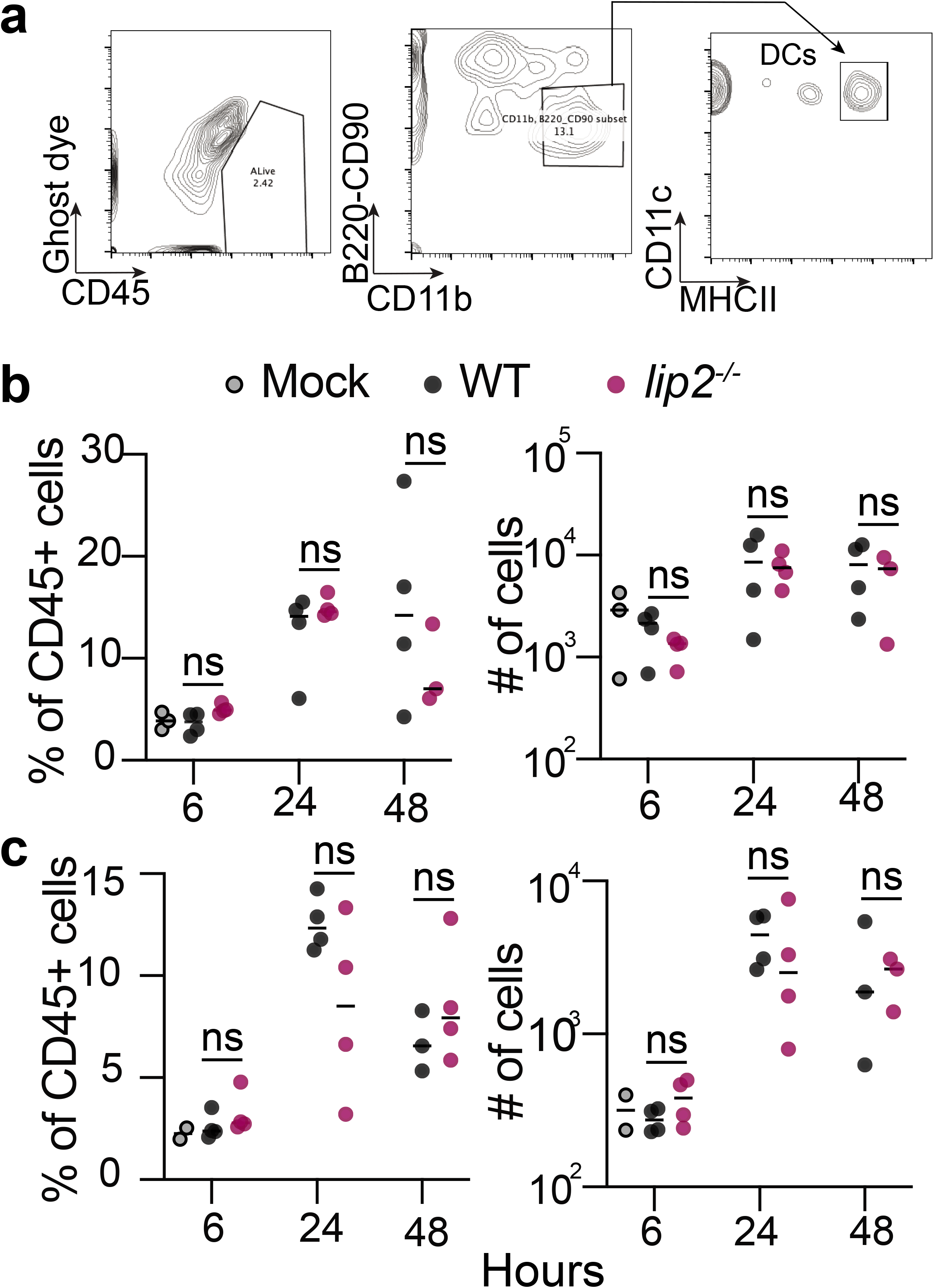
(a-b) DCs are the first responders cells. Groups of C57BL/6J mice were infected with with 1×10^6^ CFU of WT or *lip2*^−/-^ strains. Animals were euthanized after 6, 24 or 48 hours of infection. Flow cytometry analysis was performed on total renal cells and resident DCs were quantified as % of CD45 or total number of cells (fig. S3b). (c) Groups of BALB/c mice were infected with with 1×10^5^ CFU of WT or *lip2*^−/-^ strains. Animals were euthanized after 6, 24 or 48 hours of infection. Flow cytometry analysis was performed on total renal cells and resident DCs were quantified as % of CD45-positive cells or total number of cells.

**Extended Data Fig. 4.**
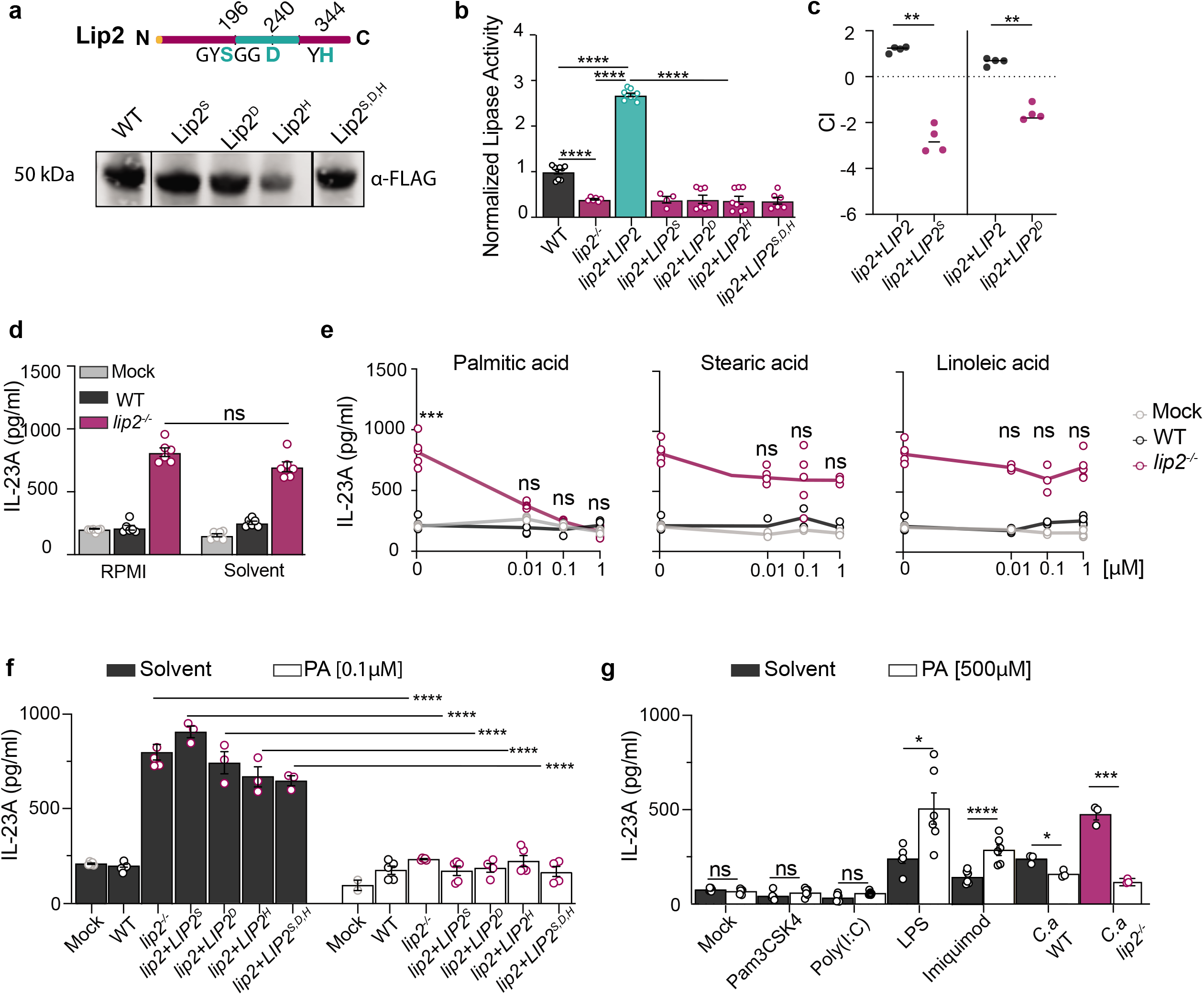
**(a)** Catalytic mutants are rightly expressed. Schematic representation of Lip2 protein domains. The green central part is the catalytic site, and the conserved amino acids are shown in green. Predicted catalytically-dead mutants were created and tagged with a 6-His-FLAG sequence (HHHHHHGGDYKDDDDK). Western blot of supernatants from catalytically-dead Lip2 mutants. (b) The catalytic site is indispensable for the in *vitro* lipase activity. Indicated strains were grown in YEPD at OD_600_=1. To assess the lipase activity, supernatant of cultures were then incubated with a a mixture of lipids for 10 min. (c) Catalytic activity of Lip2 is essential for the virulence. BALB/c mice were infected with 1×10^5^ CFU of a 1:1 mixture of *lip2*+*LIP2* and *lip2*+*LIP2^S196A^* or *lip2*+*LIP2* and *lip2*+*LIP2^D240A^*. The relative strain abundance in kidneys was determined by qPCR. Statistical significance was determined by a paired two-tailed t-test; **p < 0.01. (d-e) Palmitic acid suppresses the BMDCs activation after *lip2^−/-^* exposure. BMDCs from C57BL/ 6J mice were incubated with 0.01 μM, 0.1 μM or 1 μM of fatty acids (palmitic acid, stearic acid or linoleic acid) solubilized in chloroform (chloroform (solvent) was used as control) and co-cultured with WT, *lip2*^−/-^ strain at a MOI of 1 for two hours followed by measurement of IL-23A in cell supernatants by ELISA. Statistical significance was determined by one-way ANOVA (Turkey’s multiple comparisons test); ns: p> 0.05; *** p<0.001. (f) Lip2-catalytic point mutants show the same phenotype as *lip2^−/-^*. BMDCs from C57BL/6J mice were incubated with 0.1 μM of chloroform (solid bars) or with 0.1 μM of palmitic acid solubilized in chloroform (clear bars). BMDCs were cocultured with the indicated strains at an MOI of 1 for two hours followed by measurement of IL-23A in cell supernatants by ELISA. Statistical significance was determined by an unpaired two-tailed t-test; ns: p> 0.05; ** p<0.001; **** p<0.0001. (g) The dampening effect of palmitic acid is specific to the *lip2^−/-^* stimulus. BMDCs from C57BL/6 mice were incubated with 0.1 μM of chloroform (solid bars) or with 500 μM of palmitic acid solubilized in chloroform (clear bars). BMDCs were stimulated for 24 hours with TLR ligands (Pam3CSK4: TLR1/2; Poly(I:C): TLR3; LPS: TLR4; Imiquimod: TLR7/8) or cocultured with WT, *lip2*^−/-^ strains at an MOI of 1 followed by measurement of IL-23A in cell supernatants by ELISA. Statistical significance was determined by an unpaired two-tailed t-test; ns: p> 0.05; * p<0.05; ** p<0.001; *** p<0.001.

## Supplementary Materials

### Methods

#### Yeast manipulations

Yeast strains and primers used for this study are described in Table S1, and Table S2. All media were prepared as in ^1^. Growth assays were performed with strains that were freshly streaked from frozen glycerol stocks onto YPD agar and incubated for two days at 30°C. A single colony was suspended in sterile distilled water to an optical density at 600 nm (OD_600_) of 1 and diluted as appropriate for the assay.

Strain construction was performed as in ^2^.

#### Studies in animals

All procedures involving animals were approved by the Institutional Animal Care and Use Committee at the University of California San Francisco and were carried out according to the National Institute of Health (NIH) guidelines for the ethical treatment of animals. Experiments were performed with 8–10 week old BALB/c (no.028) mice from Charles River Laboratories or eight week old C57BL/6J (no. 000664) WT and *il17af*^−/-^ (no. 034140) mice from Jackson Laboratories and bred in-house. Systemic infection was performed by inoculation of 1×10^5^ CFUs (or indicated inoculum) of mid-log phase yeasts into the retro bulbar sinus of animals, under isoflurane anesthesia. Animals were monitored closely and euthanized at the experimentally determined endpoints or upon development of signs of clinical morbidity (defined as BCS≤2, hunched posture, decreased motor activity), whichever was first. Kidneys were then recovered for CFU analysis, histology, RNA analysis, and/or analysis of competitive fitness. Animals infected with the *lip2*(DOX-OFF) strain were additionally treated with 0.25 mg/mL of doxycycline via drinking water beginning seven days prior to infection and continued throughout the experiment.

#### Determination of competitive index

Competitive index (CI) of strains in 1:1 competitive infections of the systemic infection model was determined as in ^3^. Strain abundance was determined by qPCR of genomic DNA prepared from CFUs recovered from the inoculum and infected kidneys, using strain-specific primers (Table S2) and a Roche LightCycler 480 instrument. Significance of observed differences was determined using the paired Student’s t-test.

#### Histology

Kidneys were fixed in 10% formalin for 24 hours and washed in 80% ethanol. Preparation of paraffin blocks and 4 μm sections as well as staining with PAS and H&E were performed by Nationwide Histology (MT).

#### Kidney dissociation and recovery of renal dendritic cells

Kidneys were minced and digested with Collagenase I (0.125mg/mL, Thermo Fisher Scientific), DNAse I (0.2 mg/mL, Millipore) in RPMI with 10% of FBS for 30 min at 37°C, prior to filtration through a 100 micron strainer (Thermo Fisher Scientific) and washing with RPMI + 2% FBS + 5 mM EDTA. Cells were suspended in FACS buffer (PBS, 2% FBS, 1 mM EDTA) and purified over a discontinuous gradient of 70% and 30% Percoll (Cytivia). Cells collected from the 70%-30% interface were washed in FACS buffer for further analysis.

Isolation of renal DCs: Cells recovered from the Percoll gradient were suspended in MACS buffer and incubated with CD11c-Ab beads for 15 min at 4°C. CD11c+ DCs were isolated using MS MACS columns per the manufacturer’s instructions. Quantitation was performed with a hemocytometer, and total RNA was extracted using an RNAeasy mini kit (Qiagen).

#### Flow cytometry

Cells were stained with antibodies against TCRgd (GL3), CD4 (GK1.5), B220 (RA3-6B2), CD45 (30-F11), IL-17A (TC11-18H10.1), hNGFR (ME20.4), CD90.2 (53-2.1), CD11b (M1/70) Ly6G (1A8), CD11c (N418), F4/80 (BM8), MHCII (M5/114.15.2) (from Biolegend, BD Biosciences or eBiosciences). To determine the source of IL-17A in kidneys, dissociated cells were incubated for two hours in MACS buffer supplemented with PMA (50 ng/ml) and ionomycin (500 ng/ml) with GolgiStop (1000X). To detect intracellular IL-17A, cells were treated with BD Cytofix buffer and Perm/Wash reagent (BD Biosciences) and then stained with anti-IL-17A (C57BL/6J) or anti-hNGFR (SMART17 C57BL/6J) in Perm/wash buffer. Samples were analyzed by FACS (BD), and data were analyzed with FlowJo software (Version 10, BD, https://www.flowjo.com/).

#### RNA isolation and RT-qPCR

Isolation of total RNA from kidneys or dendritic cells was performed using TriZol or RNAeasy Mini kits per the manufacturer’s protocols. First-strand complementary DNA (cDNA) was synthesized from 1 μg of total RNA using the Superscript III cDNA Reverse Transcription Kit (Thermo Fisher Scientific). Quantitative PCR was performed with specific primers described in Table S2.

#### Preparation of BMDCs

Bone-marrow-derived dendritic cells (BMDC) were isolated from femurs and tibias of 6- to 8-week old C57BL/6J mice (WT, *tlr2−/−, tlr4-*/-, *tlr2-*/-4−/−, *clec7a-*/-). Bone marrow was eluted from bones with RPMI + 5% FBS and filtered through a cell strainer (70 μm). The cell suspension was centrifuged at 250 × g for 5 min at 4°C, and the pellet was re-suspended in 2 ml of RPMI + 5% FBS and 1 mL/mouse of ACK erythrocyte lysis buffer (Ammonium-Chloride-Potassium Lysing Buffer, Thermo Fisher Scientific). After incubation for 7 min one ice, 20 mL PBS was added before centrifugation, and pellets were washed once in RPMI. Cells were plated onto 100 mm non-tissue culture treated culture plate in complete RPMI (10% FCS, 10 μM 2-mercaptoethanol, 25 mM HEPES, Glutamax 100X, 100 U/mL penicillin, 100 μg/mL of streptomycin sulfate and 20 ng/mL GM-CSF (Prepotech)) for three days. On day three, cells were treated with ten mL of complete RPMI. On days six, eight, and ten, adherent cells were collected, washed, and plated with a Complete RPMI supplemented with 10 ng/mL of IL-4 (Gibco). Mature DCs were used on day 12.

#### In vitro activation of dendritic cells

Murine BMDCs were seeded into 24-well tissue culture-treated plates (BD) at 1×10^5^ cells per well in one ml complete RPMI (without penicillin and streptomycin) and cultured O/N at 37°C in 5% CO_2_. Activation by specific *C. albicans* strains was assessed after coincubation for two hours at MOI 1, followed by measurement of IL-23 in cell supernatants by ELISA.

Free fatty acid (FFA) supplementation. FFA (in chloroform) was added to BMDCs in culture medium to a final concentration of 1, 0.1, 0.01 μM for two hours.

Activation of DCs by TLR ligands. 5×10^5^ cells/mL were incubated in complete RPMI and treated with 100 ng/mL Pam3CSK4, 10 μg/mL Poly(I:C), 100 ng/mL LPS, or 3 μg/mL imiquimod with or without palmitic acid conjugated with BSA when mentioned. Cells were activated for 24 hours or 2 hours.

#### Measurement of cytokine production

Cytokine production was measured by enzyme-linked immunoassay using a DuoSet ELISA kit (R&D Systems) following the manufacturer’s protocol. Briefly, for detection of IL-17A and IL-23A, 200 μL of kidney homogenate was used. For *in vitro* coculture experiments with BMDCs, 100 μL of cell supernatant was used to detect IL-23A.

#### Lipase activity assay

Lipase activity was assessed with a Lipase Assay Kit (Sigma). Briefly, strains were propagated in liquid YPD to an A_600_ of 1. One mL of each culture was pelleted, and 100 μL of the supernatant was used for the assay.

#### pNp-Palmitate ester Hydrolase Activity Assay

Lipase activity was assayed spectrophotometrically using pNp-palmitate as a substrate. The reaction buffer was composed of 50 mM Tris-HCl pH 8, 1 mg/mL Arabic gum, 0.005% of Triton X-100, and 3.9 mM pNp palmitate (Sigma). 500 μL of supernatant was concentrated using concentrator tubes (Thermo Fisher Scientific) for a final volume of one ml. Samples were incubated at 30°C in the dark for 15 min, and absorbance was read at 410 nm.

#### Immunoblotting

Strains were propagated in liquid YPD at 30°C O/N. Supernatants from 10 mL of pelleted cultures were concentrated using protein concentrator tubes (Pierce), denatured in Laemmli buffer, and boiled at 100°C for 10 min. 15 μL of each sample was analyzed by SDS-PAGE and immunoblotted with anti-FLAG (Thermo Fisher Scientific).

